# From components to communities: bringing network science to clustering for genomic epidemiology

**DOI:** 10.1101/2022.08.22.504699

**Authors:** Molly Liu, Connor Chato, Art F. Y. Poon

## Abstract

Defining clusters of epidemiologically-related infections is a common problem in the surveillance of infectious disease. A popular method for generating clusters is pairwise distance clustering, which assigns pairs of sequences to the same cluster if their genetic distance falls below some threshold. The result is often represented as a network or graph of infections. A connected component is a set of interconnected nodes in a graph that are not connected to any other node. The current approach to pairwise clustering is to map clusters to the connected components of the graph. However, the distance thresholds typically used for viruses like HIV-1 tend to yield components that exclude large numbers of infections as unconnected nodes. This is problematic for public health applications of clustering, such as tracking the growth of clusters over time. We propose that this problem can be addressed with community detection, a class of clustering methods being developed in the field of network science. A community is a set of nodes that are more densely inter-connected relative to the number of connections to external nodes. Thus, a connected component may be partitioned into two or more communities. Here we describe community detection methods in the context of genetic clustering for epidemiology, demonstrate how a popular method (Markov clustering) enables us to resolve variation in transmission rates within a giant connected component of HIV-1 sequences, and identify current challenges and directions for further work.

## 1. Introduction

Identifying groups of closely-related infections is a common problem in epidemiology. The distribution of infections in space or time is often used as proxy for their epidemiological relationships. In other words, infections that were sampled in a similar location, at a similar time, or both, may share a common source. The genetic similarity of infections can be a more convenient or informative proxy than space or time, particularly for infections that can establish a persistent, chronic infection; that can remain undiagnosed as an asymptomatic infection; and/or with a relatively low rate of transmission. For instance, there is an abundance of genetic clustering studies characterizing patterns of transmission of HIV-1 (Grabowski and Redd, 2014) and hepatitis C virus (Lamoury et al., 2015). Moreover, genetic sequences are often routinely collected as a part of public health surveillance and the clinical management of infections.

There is now an extensive literature on the use of ‘molecular’ or ‘genetic’ clusters to characterize patterns of transmission in a population (Hassan et al., 2017; Poon, 2016). Clustering on the basis of the pairwise distances among sequences (Aldous et al., 2012; Balfe et al., 1990), as measured by a genetic distance (*d*) is especially popular in part because these distances can be relatively fast to compute. Moreover, pairwise distances are immutable quantities; unlike phylogenies, they do not change with the addition of sequences to the database. Any pair of sequences that have a distance below some threshold are assigned to the same cluster. We can describe this process more formally as follows: consider a complete graph *G* = (*V, E*) where each vertex *v* ∈ *V* represents a sequence or an individual infection. Every edge *e*(*v, u*) ∈ *E* between vertices *v, u* ∈ *V* is weighted by the genetic distance between the respective sequences, *d*(*v, u*). Applying a distance threshold *d*_max_ yields a subgraph of *G* that retains the full set of vertices and a reduced set of edges, *G*′ = (*V, E*′), where *E*′ = {*e*(*v, u*) ∈ *E* : *d*(*v, u*) ≤ *d*_max_}.

A connected component is a maximal subgraph *G*_*c*_ = (*V*_*c*_, *E*_*c*_) of *G* such that any vertex *v* ∈ *V*_*c*_ can be reached from any other vertex *u* ∈ *V*_*c*_ through a path of edges in *E*_*c*_ ⊆ *E*. Any given *G*_*c*_cannot be contained within a larger connected component. Although it is seldom stated explicitly, studies that use pairwise genetic clustering almost always define clusters as connected components of at least two or more vertices. Thus, even though a single vertex is considered a component in graph theory, a single infection is generally not interpreted as a cluster of size one. Indeed, these ‘non-clustered’ infections are often excluded from visualizations of the connected components. This separation of infections into clustered and non-clustered categories is frequently used as a surrogate binary variable to assess potential transmission risk factors through logistic regression (Aldous et al., 2012; Poon et al., 2015; Ragonnet-Cronin et al., 2019). The size and composition of the connected components is determined by the distance threshold. With increasing values of *d*_max_, the vertices gradually coalesce into one giant connected component. Conversely, as *d*_max_ approaches zero, each vertex becomes isolated into its own component. Thus, clustering studies employ intermediate thresholds that yield a number of connected components of moderate size. This also tends to result in a substantial number of unclustered infections.

The cross-sectional and prospective analysis of genetic clusters is a rapidly developing area of molecular epidemiology. For instance, several studies have developed models to predict the addition of newly diagnosed infections to pre-existing clusters in a population database (Ragonnet-Cronin et al., 2016a; Wertheim et al., 2018). This predictive modeling also provides a statistical basis for optimizing *d*_max_ to given population (Chato et al., 2020). In this previous work, we also observed that a substantial fraction (*>* 50%) of new infections did not join any clusters at typical distance thresholds, making them impossible to predict.

Our postulate is that the conventional practice of defining clusters from connected components is a limiting and unnecessary constraint on this application of genomic epidemiology. Specifically, there are several studies in network science that have developed algorithms that can further partition connected components into smaller clusters (Fortunato, 2010; Karrer and Newman, 2011; Leskovec et al., 2009). These are known as community detection methods. For example, the Louvain algorithm (Blondel et al., 2008) employs a ‘bottom-up’ heuristic to search for the assignment of vertices to clusters that maximizes the modularity of the graph. Modularity is a statistic that compares the observed number of edges within clusters to a random graph (Newman, 2006). Community detection methods are predominantly associated with the analysis of large social networks (Bedi and Sharma, 2016), particularly in relation to social media (Papadopoulos et al., 2012). However, they have also been applied to biological clustering problems, *e*.*g*., predicting protein function from sequence homology (Enright et al., 2002), protein interaction networks (Gulbahce and Lehmann, 2008), and gene expression networks (Treviño III et al., 2012). In sum, this abundant literature on community detection represents an untapped resource for improving applications of genetic clustering for infectious disease epidemiology.

## 2. Example application to HIV-1 sequences

To demonstrate the use of community detection for genetic clustering, we obtained 2,915 anonymized HIV-1 *pol* sequences from GenBank (accession numbers MH352627-MH355541). These sequences were used in a retrospective study of HIV-1 transmission patterns among people attending the Vanderbilt Comprehensive Care Clinic in middle Tennessee, USA (Dennis et al., 2018). We generated a multiple sequence alignment using MAFFT version 7.3.10 (Katoh and Standley, 2013) and used the program TN93 (https://github.com/veg/tn93) to calculate the pairwise genetic distances using the Tamura and Nei (1993) formula. The resulting graphs at thresholds of *d*_max_ = 0.015 and 0.03 are displayed in Figure 1. At the 1.5% threshold used in the original study, only 65 (40.1%) of 162 ‘new’ infections sampled in the last year of the study were connected to clusters of sequences sampled prior to 2015. Increasing the threshold to 3.0% augments this number from 62 to 118 (72.8%). However, this also causes the graph to coalesce around a giant connected component of 1,752 infections. The number of connected components of size 2 or greater decreases from 253 to 73.

**Figure 1:**
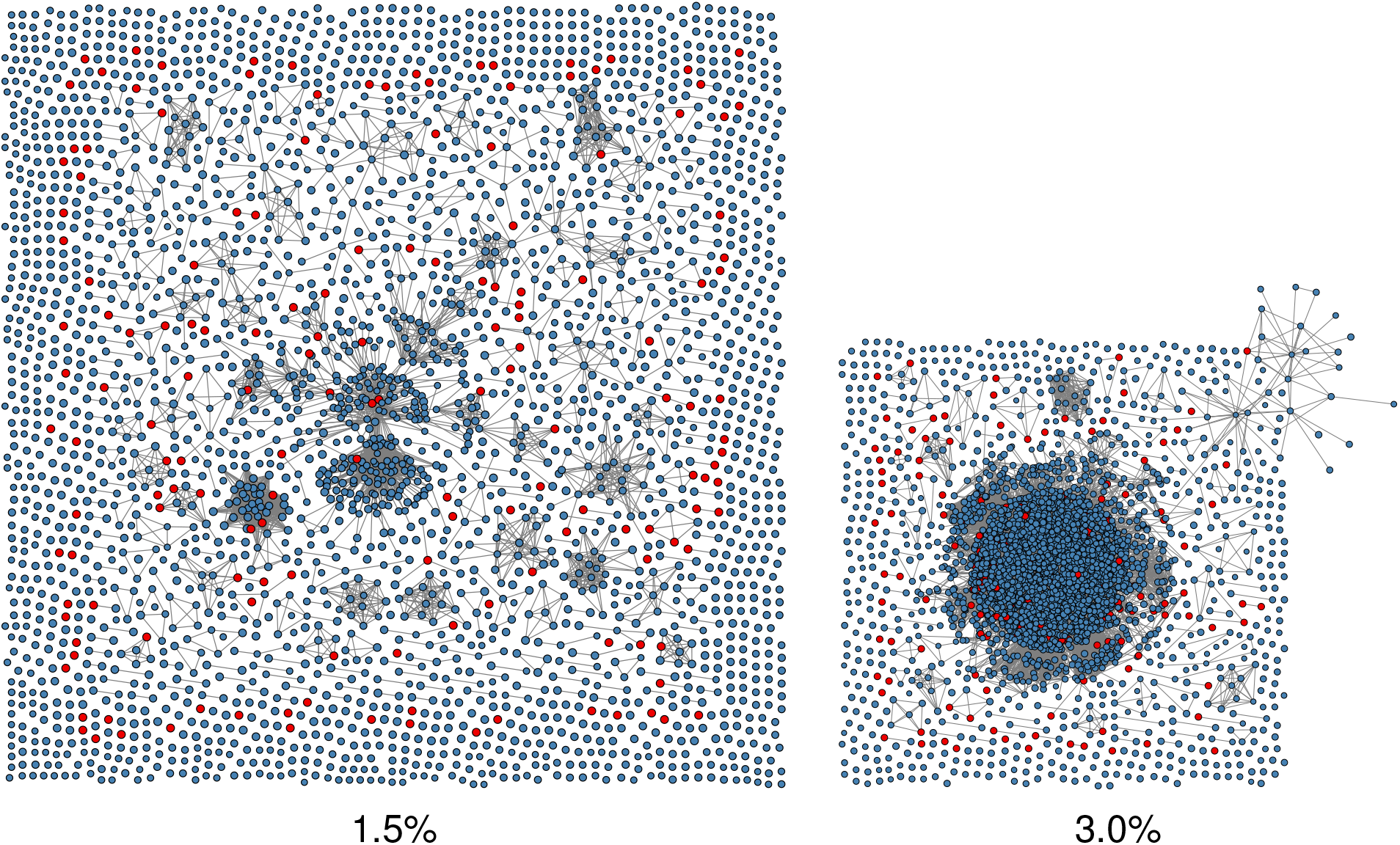
Visualizations of the graphs generated by applying thresholds of *d*_max_ = 0.015 (left) and *d*_max_ = 0.03 (right) to the pairwise distance matrix for *n* = 2, 915 HIV-1 sequences from Dennis et al. (2018). The graph layouts were generated using the ‘neato’ algorithm in GraphViz (Ellson et al., 2001). Each point represents an HIV-1 infection, with its area scaled in proportion to its year of sampling. Points are coloured red if the infection was sampled in the most recent year of the study (2015), and blue otherwise. The ‘burst’ from the upper-right corner of the 3.0% graph is due to having an insufficient number of unconnected vertices to fill the space around the connected components in this force-directed layout.

Next, we used the Poisson regression method that we previously developed (Chato et al., 2020) to determine the optimal *d*_max_ threshold for these data. The underlying concept is that the optimal threshold should yield a distribution of new infections among connected components (as clusters) that we can predict the most accurately, based on measurable characteristics of those clusters. This is quantified by the difference in the Akaike information criteria (AIC) of two Poisson regression models. The null model uses only the size of a cluster as a predictor variable, which is equivalent to assuming that every infection has the same probability of being the most closely related to a new infection (at a distance below *d*_max_). An alternate model incorporates additional predictor variables, in this case the mean time since sampling for infections in a cluster (Chato et al., 2020). We calculated the AIC of both models under a range of thresholds to yield a profile. In short, the ΔAIC for connected components was minimized at *d*_max_ = 0.0134 (Figure 2), which was fairly similar to the threshold used in the original study (0.015).

**Figure 2:**
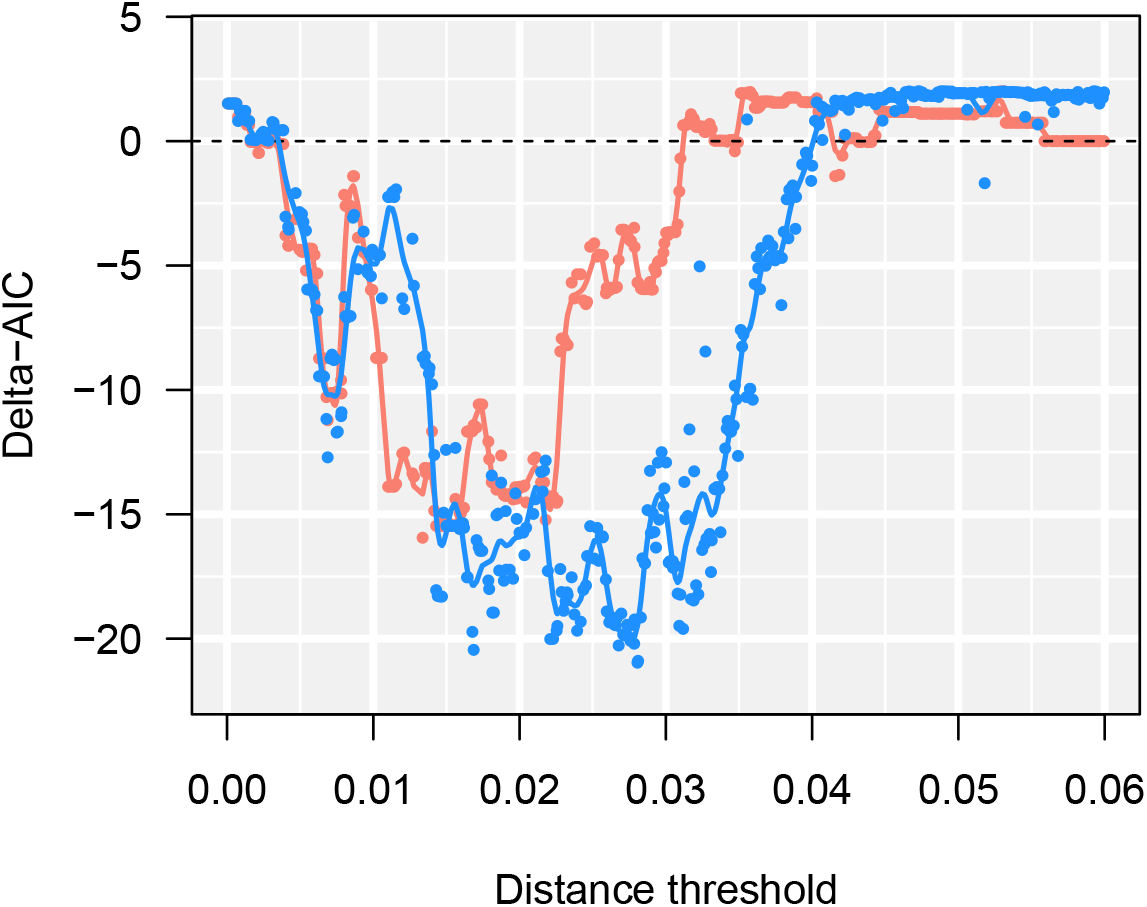
ΔAIC profiles for connected component (red) and Markov clustering (MCL, blue) methods under a range of Tamura-Nei (TN93) distance thresholds. More negative ΔAIC values indicate less information loss when incorporating additional predictor variables into a Poisson regression of new infections among clusters (Chato et al., 2020). Each point represents one of 420 parameter combinations, specifically the distance threshold (*d*_max_) and the expansion (*k*) and inflation (*r*) parameters of the MCL method. Solid lines correspond to cubic smoothing splines fit to each set of points.

Finally, we applied a community detection method known as the Markov cluster algorithm (Van Dongen, 2008) to the graphs obtained under varying thresholds, using the implementation of this method in the R package MCL (https://CRAN.R-project.org/package=MCL). MCL acts on a transition matrix (*P*) derived from the graph. In our case, we start from the adjacency matrix (*A*) of the undirected graph, where: *A*_*ij*_ = 1 if there exists an edge between vertices *i* and *j*, and 0 otherwise; *A*_*ii*_ = 0; and *A*_*ij*_ = *A*_*ji*_ ∀ *i* ≠ *j*. To derive *P* from *A*, we normalize the entries so that each column sums to 1, *i*.*e*., *P*_*ij*_ = *A*_*ij*_*/*∑_*k*_ *A*_*kj*_. Next, two different matrix operations are iteratively applied to *P*. The inflation operation takes the *r*^th^ Hadamard (entry-wise) power of *P*, such that 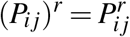, and then rescales the result so that its columns each sum to 1. The expansion operation takes the *k*^th^ power of *P* by matrix multiplication; for example, *P*^*k*^ = *PP* for *k* = 2. These operations are analogous to simulating a random diffusion process through the graph (Van Dongen, 2008). This iterative algorithm is applied until it converges to an equilibrium state where the matrices before and after operations are identical, or up to a maximum number of iterations.

We used Latin hypercube sampling to generate a uniform sample of 500 points in the space of all three parameters over the respective continuous ranges: 0 ≤ *d*_max_ ≤ 0.6; 2 ≤ *k* ≤ 25; and 2 ≤ *r* ≤ 25). Out of these MCL analyses, 80 (16%) failed to converge to an equilibrium matrix after 100 iterations. These failures tended to be associated with *d*_max_ *<* 0.025 or *d*_max_ *>* 0.04. We repeated the Poisson regression analysis on clusters produced by the MCL method to generate a ΔAIC profile with respect to *d*_max_ (Figure 2). To minimize the effect of varying *k* and *r* on estimating the optimal distance threshold, we located the minimum of a cubic smoothed spline fit to these ΔAIC values, resulting in *d*_max_ = 0.0276. This turned out to be very close to the threshold associated with the parameter combination with the lowest ΔAIC, *d*_max_ = 0.028.

The most conspicuous effect of MCL is that it partitions the largest connected component, which comprises 1,860 sequences at *d*_max_ = 0.028, into 403 clusters (Figure 3A). At this threshold, the largest component grows by 73 new infections. These infections become redistributed among 25 (6.2%) of the clusters (Figure 3B). We can also see that the clusters within this largest component that accumulated one or more new infections in 2015 tended to have more recent sampling dates than inactive clusters of the same size. Thus, even though the majority of infections have become subsumed into a single giant component, we are still able to resolve the epidemiological variation among clusters of infections within this component.

**Figure 3:**
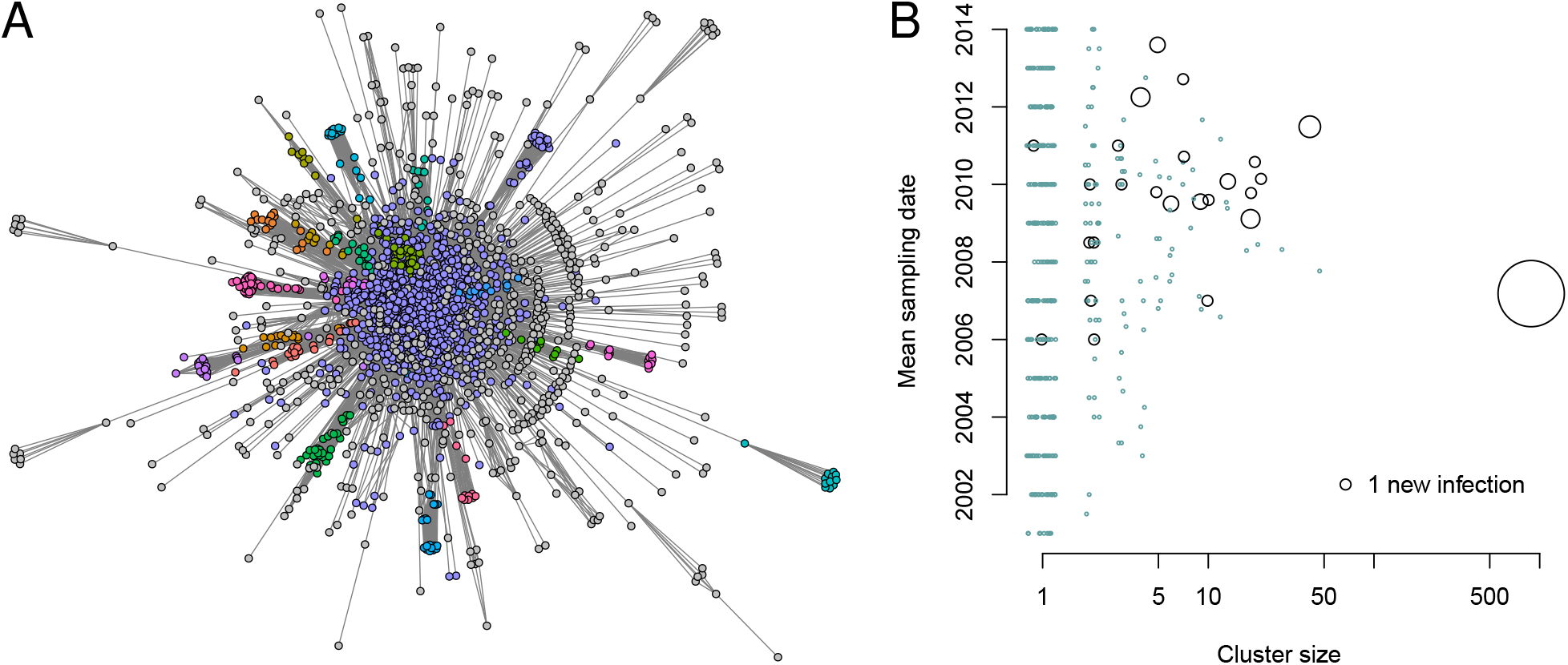
(A) Visualization of the largest connected component of HIV-1 sequences when *d*_max_ = 0.028. The vertices are coloured with respect to the 20 largest clusters as determined by the MCL algorithm, and grey otherwise. Unlike Figure 1, we used the scaleable force directed placement algorithm (*sfdp* in GraphViz; Hu, 2005) to generate a layout of this subgraph that emphasizes the separation of clusters. (B) Bubble plot summarizing the number of new cases among MCL clusters in the largest connected component. Each point represents a cluster, with its area scaled in proportion to the number of new infections added to the cluster in 2015. The smallest points, drawn in blue, represent clusters with zero new infections. We added a random ‘jitter’ to cluster size to reduce overlap.

## 3. Challenges and future directions

One basic challenge to incorporating community detection methods into the genomic epidemiology toolkit is that there are numerous and diverse methods to choose from. In addition to MCL and the Louvain algorithm, for instance, there is also stochastic blockmodeling (Karrer and Newman, 2011), convolutional neural networks (Jin et al., 2021a), and methods based on random fields (He et al., 2018); see Jin et al. (2021b) for a recent review. This may engender confusion in the field of genomic epidemiology, where many different genetic clustering methods have already been developed (*e*.*g*., Han et al., 2019; McCloskey and Poon, 2017; Villandré et al., 2018; Volz et al., 2020). Some public health agencies have already committed to a specific method of genetic clustering, such as the US Centers for Disease Control and Prevention and HIV-TRACE (Kosakovsky Pond et al., 2018; Oster et al., 2018). Thus, there would doubtless need to be some demonstrable superiority of community detection over the *status quo* for these methods to see application in the public health domain. In addition, none of these community detection methods were designed specifically for infectious disease epidemiology. Indeed we have not found any example in the literature of such methods being used to characterize viral transmission dynamics by clustering genetic sequences — at best, there is limited prior work for potential users to reference.

Incorporating community detection to a clustering analysis can introduce more parameters to calibrated by the user, in addition to distance or phylogenetic bootstrap thresholds (Hassan et al., 2017). The MCL method, for example, adds two parameters for the matrix inflation (*r*) and expansion (*k*) operations, respectively. However, we found that the ΔAIC profile that we used to optimize the distance threshold was relatively insensitive to variation in *r* and *k* (Figure S1). These results suggest that our ability to predict the distribution of new infections may be more robust to differences in community detection methods, although we have only evaluated a small number of such methods in this context. In addition, community detection methods may be too computationally complex to apply to large sequence data sets. For instance, it is not uncommon to use genetic clustering to analyze a population database comprising tens of thousands of HIV-1 sequences or more (Poon et al., 2015; Ragonnet-Cronin et al., 2016b). Fortunately, community detection methods are often designed to handle very large networks (Harenberg et al., 2014), and some have already been adapted to distributed computing environments (*e*.*g*., Azad et al., 2018).

Genetic clustering can play an important role in tracking variation in virus transmission rates in near real-time (Little et al., 2014; Poon et al., 2016). However, this emerging practice of ‘molecular surveillance’ has also raised significant concerns over ethics, consent and data privacy (Coltart et al., 2018). This is especially controversial for HIV-1, which remains a highly stigmatized infectious disease where people are criminally prosecuted for virus transmission. In this context, the phrase ‘community detection’ may be problematic, since it can be misinterpreted as an act of surveillance targeting actual communities. In many settings, communities are an important source of support, information and advocacy for people living with HIV-1 (Campbell et al., 2007). When communicating findings from applications of these methods to infectious disease epidemiology, we recommend making it clear that while community detection methods were largely developed for the analysis of social networks, they are being applied to networks where connections represent levels of genetic similarity between infections — not social links. Although networks are being used in both contexts, they are abstractions of completely different sets of relationships.

Community detection methods may be especially well-suited for pathogens with a higher transmission rate than HIV-1, such as SARS-CoV-2. When the rate of transmission exceeds the rate of molecular evolution, there is a low probability that an infection transmitted to the next host will have accumulated one or more mutations. Consequently, the distribution of pairwise distances will be shifted towards zero. In the case of SARS-CoV-2 genome sequences, setting the pairwise distance threshold to the equivalent of two nucleotide substitutions (about 6.7 × 10^−5^ expected substitutions per site) or more tends to result in giant connected components. Even at the lowest possible threshold of one mutation, we have found that pairwise distance clustering of SARS-CoV-2 genome sequences tends to yield enormous, densely-connected components, making it difficult to identify associations between individual- and group-level characteristics and transmission patterns. Thus, community detection may provide an important mechanism enabling investigators to resolve transmission patterns from genetic sequences for a much wider range of viruses than HIV-1 and hepatitis C virus.

## Author contributions

The study was conceived and designed by AFYP. ML and AFYP curated the data. ML implemented the statistical and computational methods to analyze the data. CC provided analytical tools. AFYP and ML drafted the manuscript and generated data visualizations. AFYP, ML and CC critically reviewed and revised the manuscript.

## Supplementary Figures

**Figure S1:**
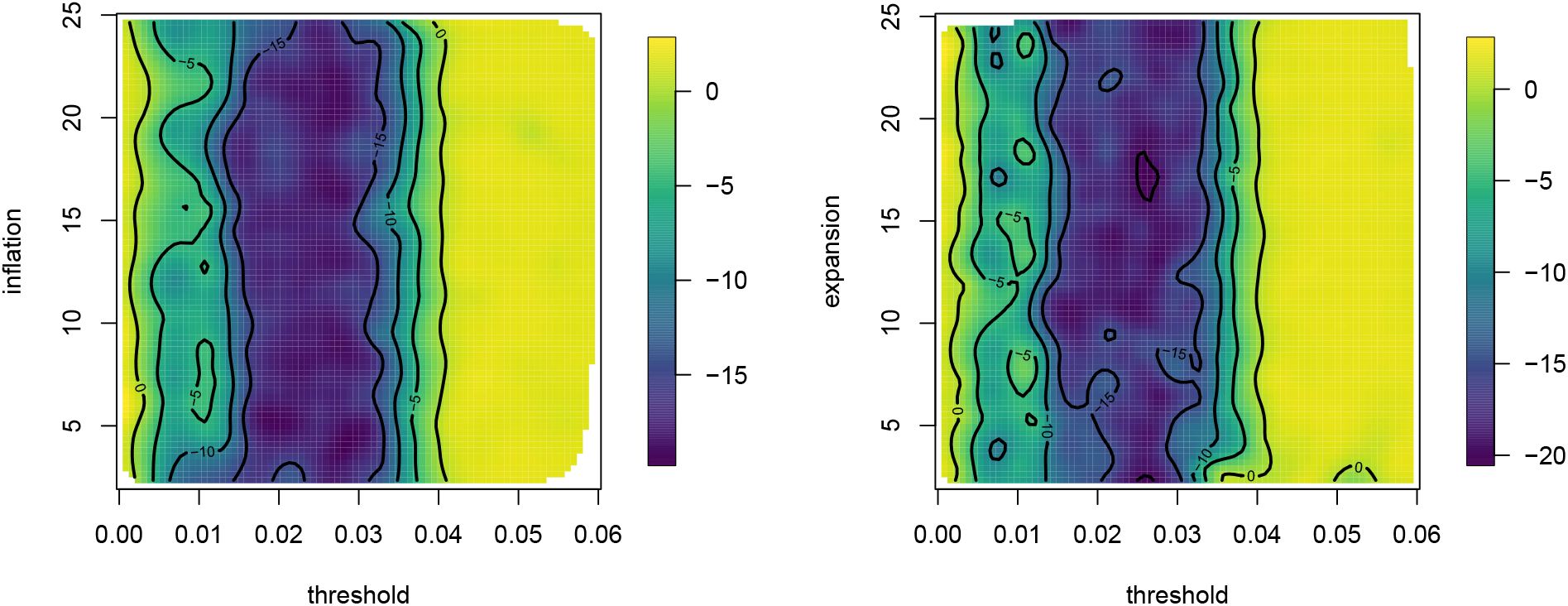
Filled contour plots summarizing the response of ΔAIC to variation in the Markov clustering parameters for inflation (top) and expansion (bottom) matrix operations, over a range of Tamura-Nei (TN93, horizontal axis) distance thresholds. Essentially, they are ‘stretched’ versions of Figure 2. Darkest colours correspond to the most negative values of ΔAIC. These plots were generated by fitting a thin plate spline regression, using the Tps function in the R package *fields* (Nychka et al., 2021), to *n* = 614 ΔAIC values generated from our Poisson regression analyses of the HIV-1 sequence data.

